# The effect of variant interference on *de novo* assembly for viral deep sequencing

**DOI:** 10.1101/815480

**Authors:** Christina J. Castro, Rachel L. Marine, Edward Ramos, Terry Fei Fan Ng

**Affiliations:** Division of Viral Diseases, National Center for Immunization and Respiratory Diseases, Centers for Disease Control and Prevention, Atlanta, Georgia, USA; Oak Ridge Institute for Science and Education, Oak Ridge, Tennessee, USA; General Dynamics Information Technology, Inc., contracting agency to the Office of Informatics, National Center for Immunization and Respiratory Diseases, Centers for Disease Control and Prevention, Falls Church, VA, USA

**Author notes:** To whom correspondence should be addressed: Terry Fei Fan Ng, Division of Viral Diseases, Centers for Disease Control and Prevention 1600 Clifton Rd. NE, Mailstop H17-6 Atlanta, GA 30329.

## Abstract

Viruses have high mutation rates and generally exist as a mixture of variants in biological samples. Next-generation sequencing (NGS) approach has surpassed Sanger for generating long viral sequences, yet how variants affect NGS *de novo* assembly remains largely unexplored. Our results from >15,000 simulated experiments showed that presence of variants can turn an assembly of one genome into tens to thousands of contigs. This “variant interference” (VI) is highly consistent and reproducible by ten most used *de novo* assemblers, and occurs independent of genome length, read length, and GC content. The main driver of VI is pairwise identities between viral variants. These findings were further supported by *in silico* simulations, where selective removal of minor variant reads from clinical datasets allow the “rescue” of full viral genomes from fragmented contigs. These results call for careful interpretation of contigs and contig numbers from *de novo* assembly in viral deep sequencing.

## Introduction

For many years, Sanger sequencing has been used to complement classical epidemiological and laboratory methods for investigating viral infections.^1^ As technologies have evolved, the emergence of next-generation sequencing (NGS), which drastically reduced the cost per base to generate sequence data for complete viral genomes, has allowed scientists to apply viral sequencing on a grander scale.^2^ Genomic sequencing is ideal for elucidating viral transmission pathways, characterizing emerging viruses, and locating genomic regions which are functionally important for evading the host immune system or antivirals.^3^

Genomic surveillance of viruses is particularly important in light of their rapid rate of evolution. Viruses have higher mutation rates than cellular-based taxa, with RNA viruses having mutation rates as high as 1.5 × 10^−3^ mutations per nucleotide, per genomic replication cycle.^4^ Due to this high mutation rate, it is well established that most RNA viruses exist as a swarm of quasispecies,^5^ with each quasispecies containing unique single nucleotide polymorphisms (SNPs). The presence of these variants plays a key role in viral adaptation.

Due to viruses’ rapid evolution, a single clinical sample often contains a mixture of many closely related viruses. Viral quasispecies are mainly derived from intra-host evolution, with RNA viruses such as poliovirus, human immunodeficiency virus (HIV), hepatitis C (HCV), influenza, dengue, and West Nile viruses maintaining diverse quasispecies populations within a host.^6, 7, 8, 9, 10, 11, 12, 13^ Conversely, the term “viral strains” often refers to different lineages of viruses found in separate hosts, or a co-infection of viruses in the same host due to multiple infection events. As a result, sequence divergence is usually higher when comparing viral strains compared to quasispecies. In this study, we use the term “variant” to encompass both quasispecies and strains regardless of how the variants originated in the biological samples.

Since many sequencing technologies produce reads that are significantly shorter than the target genome size, a process to construct contigs, scaffolds, and full-length genomes is needed. Reference-mapping and *de novo* assembly are the two primary bioinformatic strategies for genome assembly. Reference-mapping requires a closely-related genome as input to align reads, while *de novo* assembly generates contigs without the use of a reference genome, and therefore is the most suitable strategy for analyzing underexplored taxa^14^ or for viruses with high mutation and/or recombination rates.

In this study, we first examined how often NGS and *de novo* assembly were applied in viral sequencing in GenBank Nucleotide entries (www.ncbi.nlm.nih.gov/nucleotide/). Then we investigated how the presence of variants affected assembly results - simulated and clinical NGS datasets were analyzed using multiple assembly programs to explore the effects of genome variant relatedness, read length, and genome length on the resulting contig distribution.

## Results

### The rise of NGS and *de novo* assembler use in GenBank viral sequences

GenBank viral entries from 1982-2017 were collected and analyzed, with extensive analyses performed to evaluate technologies and bioinformatics programs cited in records deposited between 2011 and 2017. Through 2017, there were over 2.3 million viral entries in GenBank; however, over 70% (1.7 million) do not specify a sequencing technology [Supplement Table S1] due to the looser data requirement in earlier years. When looking at recently deposited records (2014-2017), the Illumina sequencing platform was the most common NGS platform used for viral sequencing, with about a 2-fold increase over the next most popular NGS platform [Figure 1d & e]. When long sequences (≥2,000 nt) are considered, NGS technologies surpassed Sanger in 2017 as the dominant strategy for sequencing, comprising 53.8% (14,653/27,217) of entries compared to 46.2% of entries (12,564/27,217) for Sanger [Figure 1f and Supplement Table S2].

Hybrid sequencing approaches, where researchers use more than one sequencing technology to generate complete viral sequences, have also become more common over the past several years. The most common combination observed was 454 and Sanger (18,002 entries), likely due to the early emergence of the 454 technology compared to other NGS platforms [Figure 1c and Supplement Table S3]. However, combining Illumina with various other sequencing platforms is quite commonplace (>10,000 entries).

**Figure 1.**
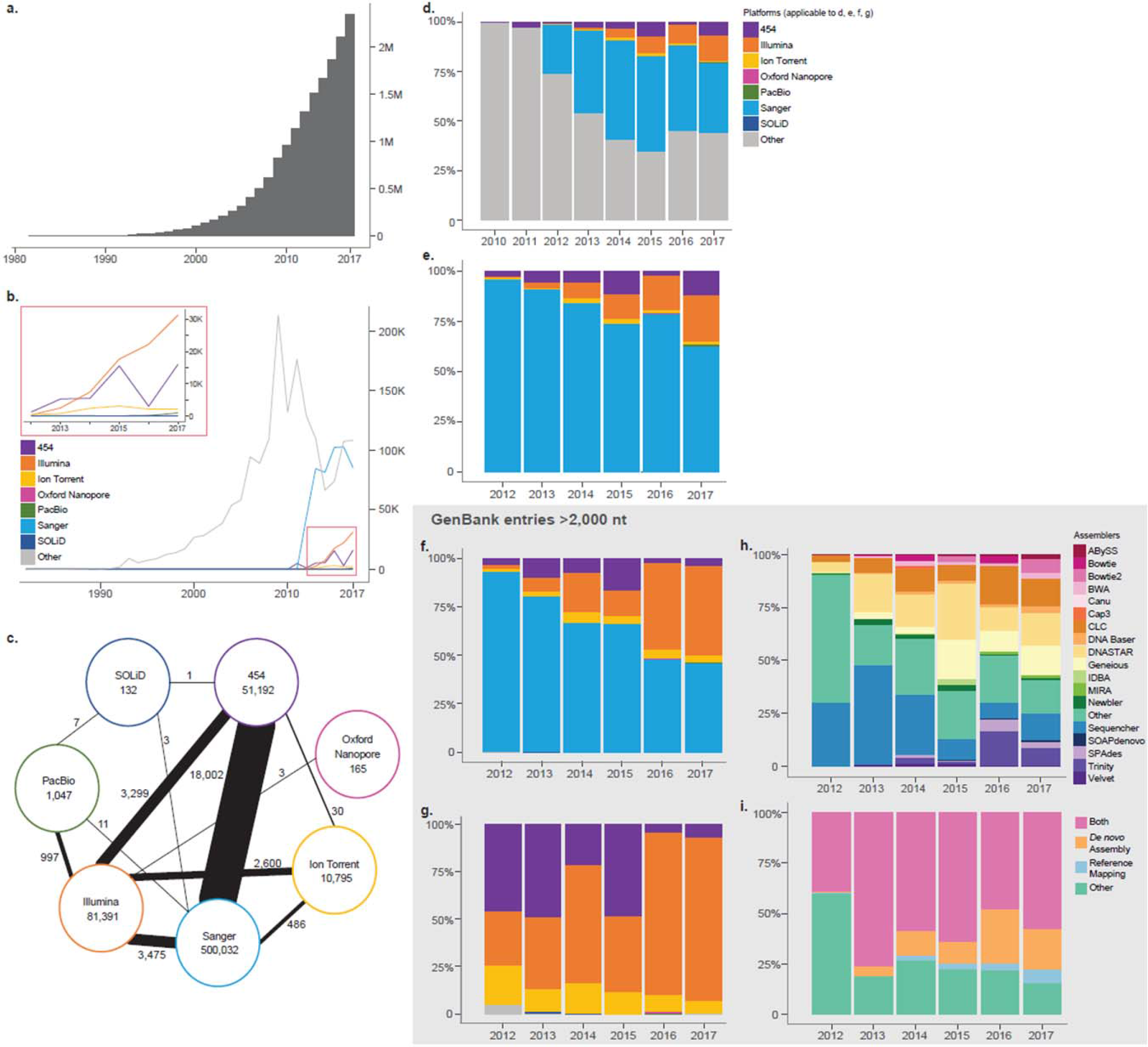
Trends and patterns of sequencing technology and assembly methods of viral entries in the GenBank database. **(a)** Cumulative frequency histogram of all viral entries in GenBank from Jan. 1, 1982 through Dec. 31, 2017 (total=2,338,775 entries). **(b)** Count of all viral entries with at least one *Sequencing Technology* documented for the years 1982-2017. For panels (b) and (d), the “Other” category denotes entries with the *Sequencing Technology* field omitted or mis-assigned. **(c)** Relationship between viral entries listing one or two *Sequencing Technologies* during 1982–2017. The number inside the circle indicates viral entries with only one *Sequencing Technology* listed; the number adjacent to the line indicates entries combining two *Sequencing Technologies*. The thicker the connection line, the stronger the relationship. **(d and e)** Percentage ratio graph of all viral entries with *Sequencing Technology* documented for the years 2010–2017, with (d) and without (e) the *Other* category. The majority of entries in earlier years include omissions classified under the *Other* category, which is detailed in Supplement Table S1. **(f)** Percentage ratio graph of viral entries with length greater than 2000 nt that have been documented with one of the seven *Sequencing Technologies* for the years 2012–2017. The seven technologies includes Sanger (n=1) and NGS technologies (n=6). **(g)** Percentage ratio graph of viral entries with length greater than 2000 nt and that have been documented with one of the six NGS as the *Sequencing Technology* for the years 2012–2017. Compared to panel (f), Sanger is excluded in this graph. **(h)** Assembly method of viral entries greater than 2000 nt, showing percentage ratio graph of entries with at least one *Assembly Method*. For (h) and (i), the *Other* category describes assembly methods outside of the 18 most popular programs investigated. **(i)** Reclassification of panel (h) by the nature of the assembly methods. The programs can be grouped into *de novo* assembler, reference-mapping assembler, and software that can perform both.

*De novo* assembly programs (ABySS, BWA, Canu, Cap3, IDBA, MIRA, Newbler, SOAPdenovo, SPAdes, Trinity, and Velvet) have increased from less than 1% of viral sequence entries in 2012, to 20% of all viral sequence entries in 2017 [Figure 1h & i]. A similar increase was observed for reference-mapping programs (i.e., Bowtie and Bowtie2), from 0.03% in 2012 to 6.5% in 2017. Multifunctional programs (Suppl. Information) that offer both assembly options were the most common programs cited for the years 2013-2017, but since the exact sequence assembly strategy used for these records is unknown, the contributions of *de novo* assembly are likely underestimated. An expanded summary of the sequencing technologies and assembly approaches used for viral GenBank records is available in Supplement Tables S1–S6.

### Effect of variant assembly using popular *de novo* assemblers

After establishing the growing use of NGS technologies for viral sequencing, we next focused on understanding how the presence of viral variants may influence *de novo* assembly output. We generated 247 simulated viral NGS datasets representing a continuum of pairwise identity (PID) between two viral variants, from 75% PID (one nucleotide difference every 4 nucleotides), to 99.6% PID (one nucleotide difference every 250 nucleotides) [Figure 2]. For Experiment 1, these datasets were assembled using 10 of the most used *de novo* assembly programs [Figure 2 and Supplement Figure S1a] to evaluate their ability to assemble the two variants into their own respective contigs as the PID between the variants increases.

**Figure 2.**
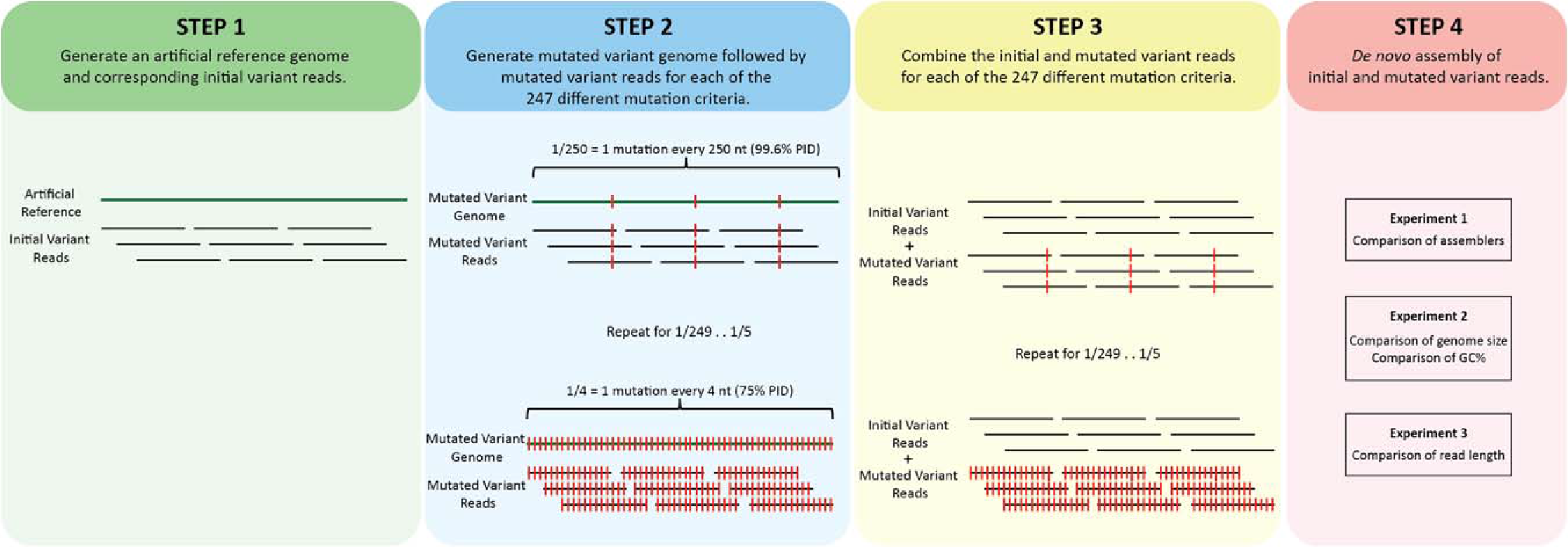
Workflow diagram of the investigation of variant simulated NGS reads through *de novo* assembly. First, in step 1, an artificial reference genome and corresponding initial variant reads were created with varying constraints such as genome length, GC content, read length, and assemblers, according to the experiment types as detailed in Supplement Figure S1. In the second step, an artificial mutated variant genome was created. The process is repeated to generate 247 different mutated variants with controlled mutation parameters— starting with 1 mutation every 4 nucleotides (75% PID) and ending with a mutated variant with 1 mutation in every 250 nucleotides (99.6% PID). Mutated variant reads are also generated for each of the mutation parameters. In the third and fourth steps, the initial and mutated variants were then combined and used as input for *de novo* assembly for the three experiments, as detailed in Supplement Figure S1.

One key observation is that the assembly result can change from two (correct) contigs to many (unresolvable) contigs simply by having variant reads; the presence of viral variants affected the contig assembly output of all 10 assemblers tested. The output of the SPAdes, MetaSPAdes, ABySS, Cap3, and IDBA assemblers shared a few commonalities, demonstrated by a conceptual model in Figure 3A. First, below a certain PID, when viral variants have enough distinct nucleotides to resolve the two variant contigs, the *de novo* assemblers produced two contigs correctly [Figure 3]. We refer to this as “variant distinction” (VD), with the highest pairwise identity where this occurs as the VD threshold. Above this threshold, the assemblers produced tens to thousands of contigs [Figure 3], a phenomenon we define as “variant interference” (VI). As PID between the variants continue to increase, the *de novo* assemblers can no longer distinguish between the variants and assembled all the reads into a single contig, a phenomenon we define as “variant singularity” (VS). [Figure 3]. The lowest pairwise identity where a single contig is assembled is the VS threshold.

**Figure 3.**
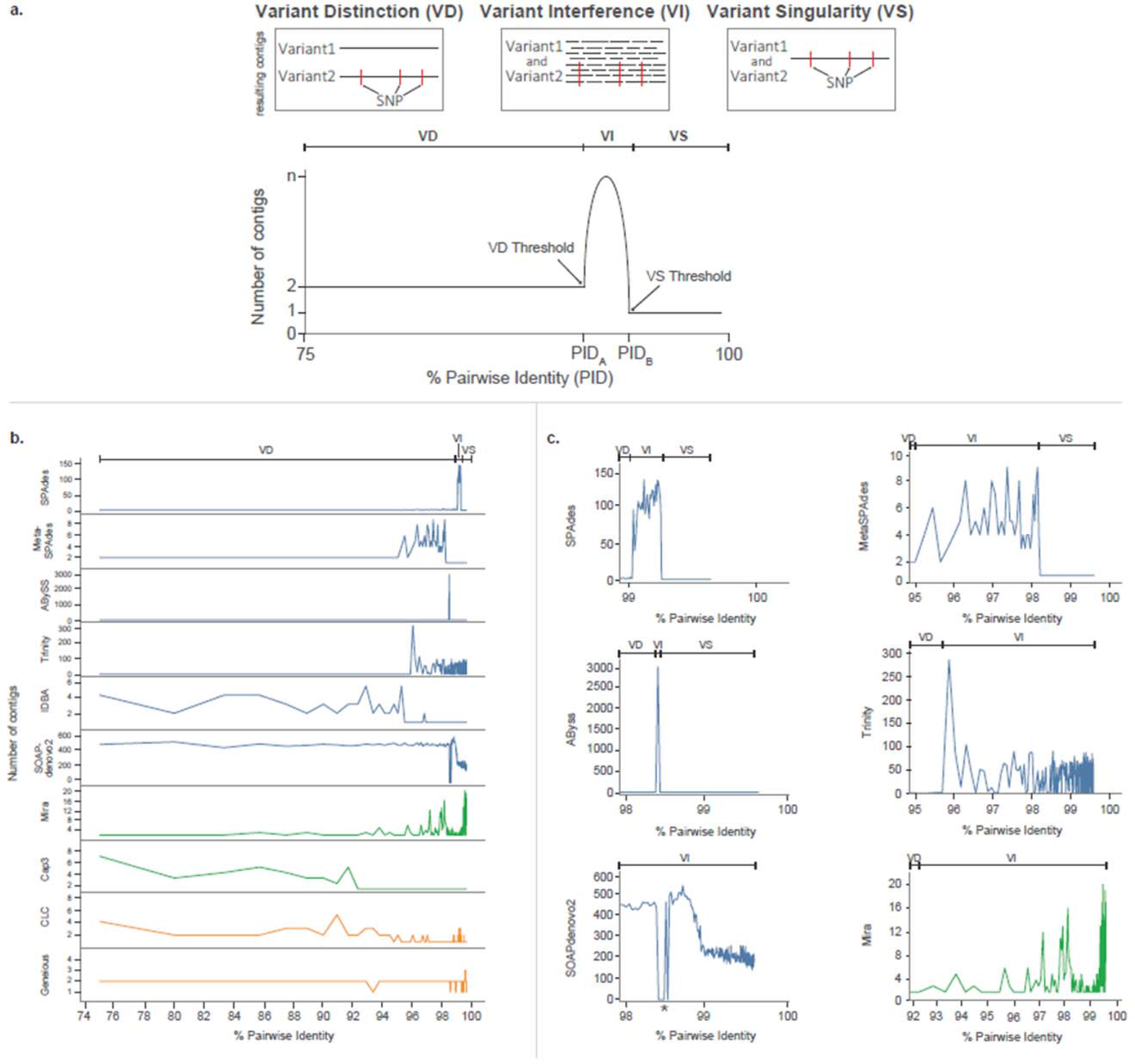
The number of contigs generated by different *de novo* assemblers using simulated data containing variants differed with a range of percentage identities (PID). Blue denotes de Bruijn graph assemblers (DBG); green denotes overlap-layout-consensus assemblers (OLC); orange denotes commercialized proprietary algorithms. Variant distinction, VD; variant interference, VI; variant singularity, VS. *For SOAPdenovo2, several data points returned zero contigs due to a well-documented segmentation fault error. **(a) Schematic diagram depicting concepts of the VD, VI, and VS, and their relationship to PID. (b) Comparison of output from 10 different assemblers.** The number of contigs produced by each *de novo* assemblers at different variant PID ranges (75%–99.6%) were shown. **(c) Close-up of PID ranges where variant interference is the most apparent.** The y-axis denotes the number of contigs.

Slight differences in the variant interference patterns (relative to the canonical variant interference model) were observed for the 10 assemblers investigated. VD was observed for SPAdes, MetaSPAdes, and ABySS assemblers. While it was not observed with Cap3 and IDBA with the current simulated data parameters, we speculate that VD may occur at a lower PID level for these assemblers than tested in this study. The PID range where VI was observed was distinct for each *de novo* assembler [Figure 3]. During VI, SPAdes produced as many as 134 contigs and ABySS produced 3,076 contigs, while MetaSPAdes, Cap3, and IDBA produced up to 10.

A different pattern was observed for Mira, Trinity, and SOAPdenovo2 assemblers. The average number of contigs generated by Mira, Trinity, and SOAPdenovo2 was 5, 36, and 283, respectively across all variant PIDs from 75%–99.96%. Specifically, Mira and Trinity generated fewer contigs at low PID, but produced many contigs when the two variants reach 97.1% PID and 96.0% PID, respectively. For SOAPdenovo2, a larger number of contigs were produced regardless of the PID. This indicates that these assemblers generally have major challenges producing a single genome; this has been observed in previous studies comparing assembly performance.^15^

Finally, Geneious and CLC were the least affected by VI in the simulated datasets tested, returning only 1–5 contigs for all pairwise identities. CLC’s assembly algorithm primarily returned a single contig over the range of PIDs tested (218/247 simulations; 88.3%), thus favoring VS. In comparison, Geneious predominantly distinguished the two variants (234/247 simulations; 94.7%), favoring VD.

### Effect of GC content and genome length on variant assembly

For Experiment 2, we focused our study on evaluating whether VI observed in SPAdes *de novo* assembly is influenced by the GC content or genome length of the pathogen. Two datasets were used for the evaluation: reads generated from four artificial genomes ranging in length from 2 Kb to 1 Mb, as well as from genome sequences of poliovirus (NC_002058; 7,440 nt in length) and coronavirus (NC_002645; 27,317 nt in length). No discernable correlation was observed between the GC content of variant genomes and the degree of VI for any of the simulated datasets [Supplemental Dataset S2, p < 0.0001]. Therefore, for subsequent analyses examining the effects of genome length on VI, the number of contigs at each PID level was obtained by averaging the 13 GC simulations.

Notably, no matter the genome length, SPAdes produced vastly more contigs (i.e., VI) in a constant, narrow range of PID [99%–99.21% ; Figure 4a & b]. The effect of variants on assembly was characterized by the three distinct intervals described previously: VD at lower PIDs, VI [Figure 4b], and VS at higher PIDs for all genome lengths. For example, during VS, a single contig was generated when the two variants shared ≥99.22% PID, but tens to thousands of contigs were generated at a slightly lower PID of 99.21%. This PID threshold, 99.21%, marked the drastic transition from VS to VI, whereas the transition from VI to VD (i.e., the VD threshold) occurred at 98.99% PID [Figure 4b]. A correlation was observed between genome length and the number of contigs produced during VI, where longer genomes returned proportionally more contigs as expected as total VI occurrence should increase with length [*r*^2^ = 0.967; p <0.0001 Figure 4b and 4c].

**Figure 4.**
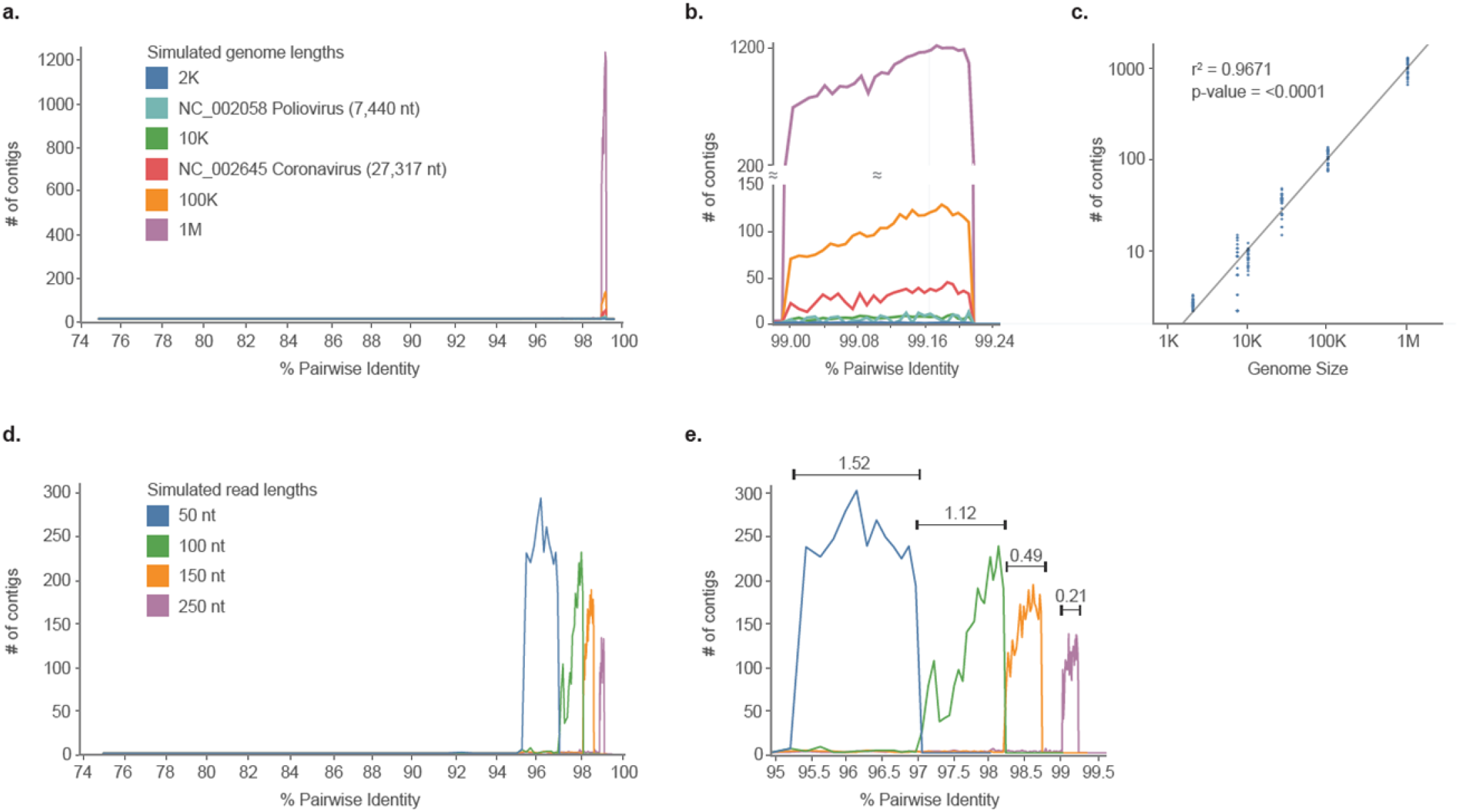
The effect of genome length and read length on *de novo* assembly of simulated variants across a range of percentage identities (PID). (a & b) Comparison of genome lengths. Six different genome lengths were assembled and the final contig counts were tallied across varying PID thresholds (75%–99.6%). For the simulated genome lengths of 2Kb, 10kb, 100Kb, and 1Mb, the average of contig number at each PID was plotted. Panel (b) shows the close-up view where interference was the most prominent. For all six genome lengths and each of the 13 iterations, VI consistently occurred in the same range of PID (99.00%–99.24%). The assembly makes a transition from VD to VI at the threshold of 99.00%, and it makes a transition from VI to VS at the threshold of 99.24%. Also, the longer the genome length, the more contigs produced during VI. **(c) The relationship between genome length and the total number of contigs produced.** Data from panel (a) were plotted on a logarithmic scale. The total number of contigs produced is significantly dependent on the genome size (r^2^=0.967; p-value<0.0001). **(d and e) The effect of read length in variant assembly with a genome size of 100K.** Simulated data with four different read lengths were created and assembled, and the final contig counts were tallied across varying PID thresholds (75%–99.6%). Panel (e) shows the close-up view where interference was the most apparent. When longer read lengths were used, the variant interference PID range was much narrower than when shorter read lengths were used to build contigs.

### Effect of read length on variant assembly

The read length of a given NGS dataset will vary depending on the sequencing platform and kits utilized to generate the data. Since read length is an important factor for *de novo* assembly success,^16^ we hypothesized that it may also influence the ability to distinguish viral variants. For Experiment 3, using SPAdes we investigated assemblies with four typical read lengths: 50, 100, 150, and 250 nt. At longer read lengths, the VD threshold occurred at higher PIDs [Figure 4d & e]. Also, with increasing read length, the width of the PID window where VI occurs gradually decreased from a 1.52% spread to a 0.21% spread [Figure 4e]. This indicates that longer reads are better for distinguishing viral variants with high PIDs.

### *In silico* experiments examining variant assembly with NGS data derived from clinical samples

For clinical samples, assembly of viral genomes is affected by multiple factors other than the presence of variants, including sequencing error rate, host background reads, depth of genome coverage, and the distribution (i.e., pattern) of genome coverage. We next utilized viral NGS data generated from four picornavirus-positive clinical samples (one coxsackievirus B5, one enterovirus A71, and two parechovirus A3) to explore VI in datasets representative of data that may be encountered during routine NGS. The NGS data for each sample was partitioned into four bins of read data: (1) total reads after quality control (**T**); (2) major variants only (**M**); (3) major and minor variants only (**Mm**); and (4) major variants and background non-viral reads only (**MB**) [Figure 5]. These binned datasets were then assembled separately using three assembly programs: SPAdes, Cap3, and Geneious. By comparing these manipulations, we aimed to test the hypothesis that minor variants directly affect the performance of assembly through VI in real clinical NGS data.

**Figure 5.**
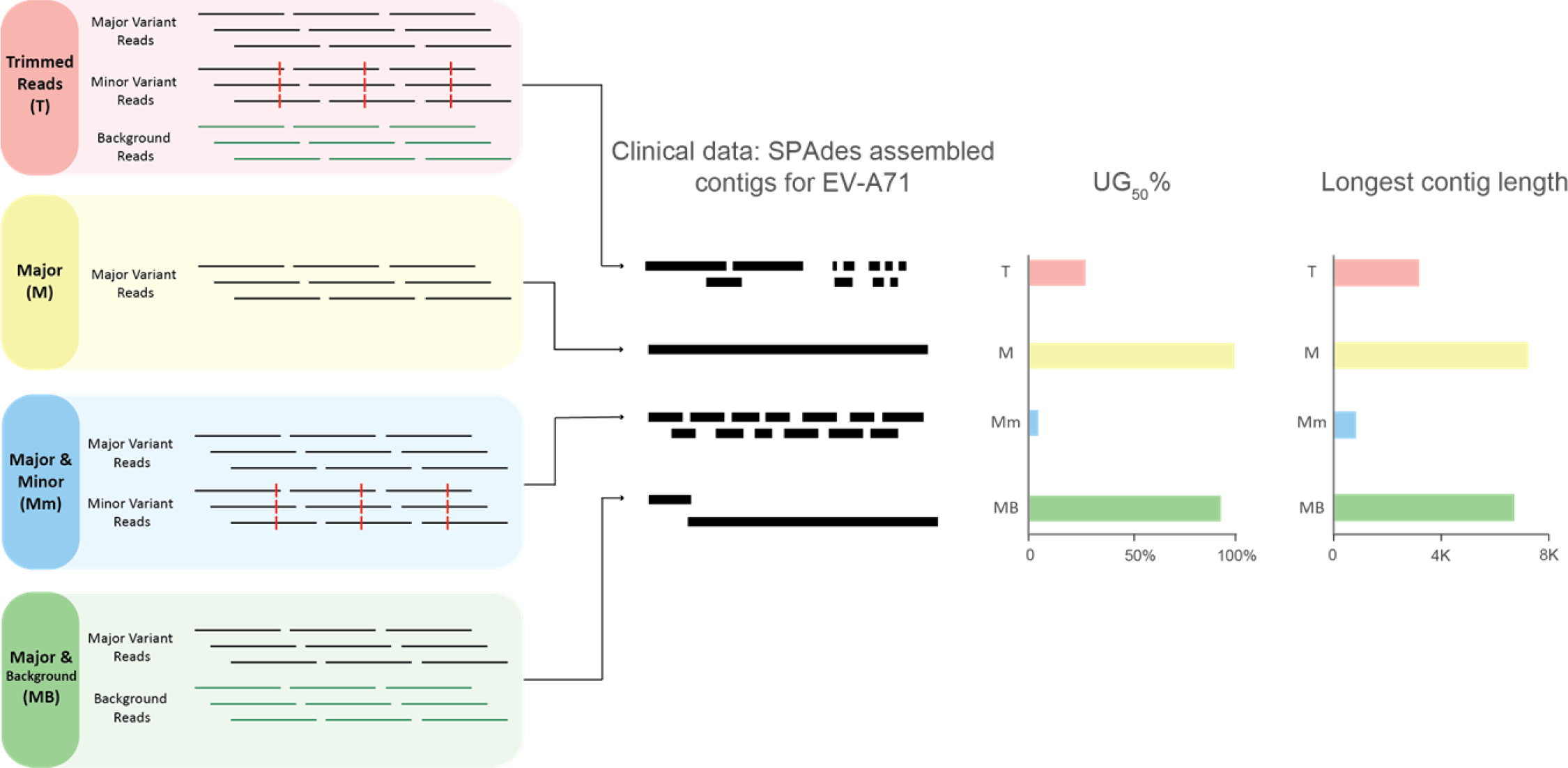
The effect of variant interference in a real dataset from a clinical sample containing enterovirus A71 (EV-A71) and its variants. Fastq reads were partitioned into four components: trimmed reads after quality control (T), major variant (M), minor variant (m), and background (B). These reads were then combined into four different experiments: T, M, Mm, and MB and assembled using SPAdes. The contig representation schematic showing the abundance and length of the generated contigs reveals the impact of variant interference on *de novo* assembly. The bar graphs show the UG_50_% metric and the length of the longest contig. UG_50_% is a percentage-based metric that estimates length of the unique, non-overlapping contigs as proportional to the length of the reference genome.^17^ Unlike N_50_, UG_50_% is suitable for comparisons across different platforms or samples/viruses. More clinical samples and viruses are analyzed similarly in Figure 6.

Even with an adequate depth of coverage for genome reconstruction, assembly of total reads (**T**) in 11/12 experiments resulted in unresolved genome construction – resulting in numerous fragmented viral contigs [Figure 6]. The only exception was one experiment where one single PeV-A3 (S1) genome was assembled using Cap3. When only reads from the major variant were assembled (**M**), full genomes were obtained for all datasets using SPAdes and Cap3, and for the CV-B5 sample using Geneious. Conversely, assembly of the read bins containing major and minor variants (**Mm**) resulted in an increased number of contigs for 9 of the 12 sample and assembly software combinations tested [Figure 6], indicating that VI due to the addition of the minor variant reads likely adversely affected the assembly. The presence of background reads with major variant reads (**MB**) did not appear to affect viral genome assembly, as the UG_50_% value, a performance metric which only considers unique, non-overlapping contigs for target viruses^17^, was similar between **M** and **MB** datasets.

**Figure 6.**
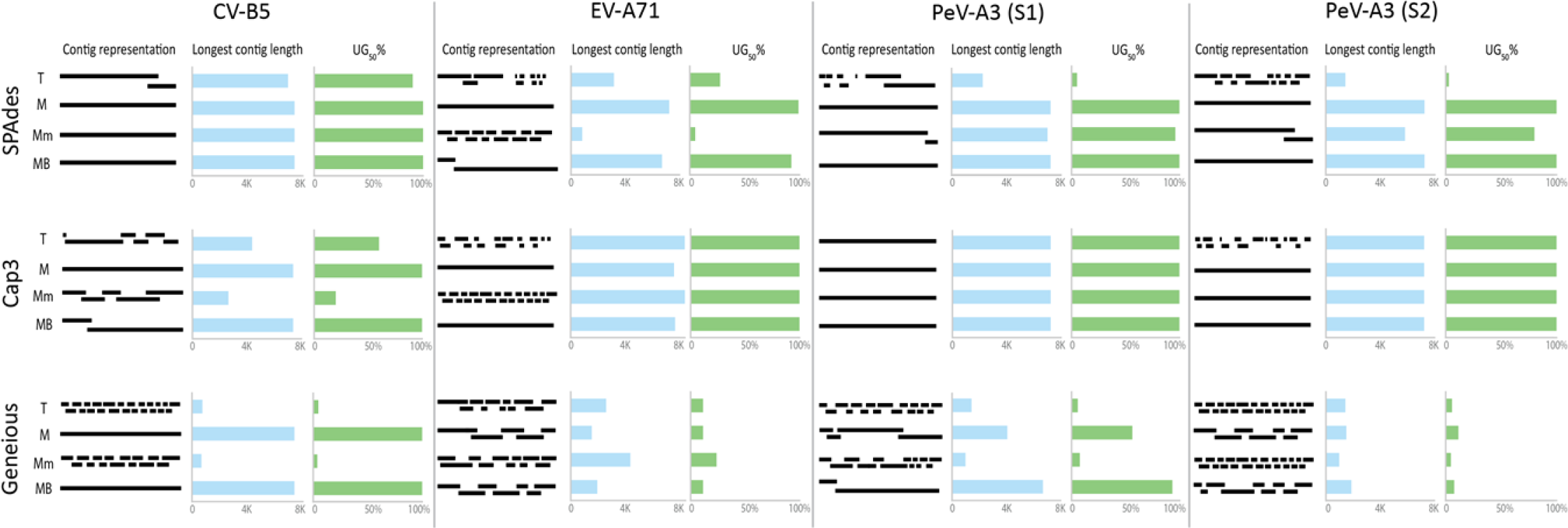
The effect of variant interference on the assembly of four clinical datasets using three assembly programs. Fastq reads were partitioned into four categories: total reads (T), major variant (M), minor variant (m), and background (B). These reads were then combined into four different categories: T, M, major and minor variants (Mm), and major variant and background (MB). Datasets were assembled using SPAdes, Cap3, and Geneious. The bar graphs show the UG_50_% metric and the length of the longest contig. Coxsackievirus B5, CV-B5; Enterovirus A71, EV-A71; Parechovirus A3 (Sample 1), PeV-A3 (S1); Parechovirus A3 (Sample 2), PeV-A3 (S2).

## Discussion

Our analysis of the GenBank quantified the decade-long expansion of NGS technologies and *de novo* assembly for viral sequencing [Figure 1]. As the number of viral sequences in public databases continues to grow, an important question that naturally arises is how well current *de novo* assembly programs perform for datasets with viral variants. Viral variants are expected in biological samples, with the number of variants and the extent of the sequence divergence between variants related to the mutation rate of the virus and the types of specimens that are being investigated. For example, samples containing rapidly evolving RNA viruses, such as poliovirus, HIV, and HCV^7, 9, 18^, environmental samples,^19^ and clinical samples from immunosuppressed individuals^20, 21^ usually harbor many variants. The ability to accurately distinguish variants is imperative to inform treatments (in the case of HIV and HCV), or determine whether a subpopulation of a more virulent variant is present.

Several experiments using simulated and clinical sample NGS data were performed to evaluate the ability of genome assembly programs to distinguish genome variants. All assemblers investigated generated fragmented assemblies when the data contained reads from two closely related variants due to “variant interference” (VI). Changes in pairwise identity (PID) as small as 0.01% between the two variants triggered an assembler to change from producing one or two contigs to producing hundreds of contigs. A quintessential example of this phenomenon was the SPAdes assembly of EV-A71 sequences during the *in silico* experiments with clinical NGS data. Assembly of major variant reads resulted in one full length contig [Figure 5], whereas assembly of datasets containing the major and minor variant reads (**Mm** and **T**) were characterized by a number of contigs, resulting in “cobwebs” of contig fragments when visualized using Bandage [Supplement Figure S2].^22^ Even though the *de novo* assembly graph linked the different contig fragments, the assembly could not differentiate the multiple routes of possible contig construction. We speculate this is the main reason why VI occurs in the context of de Bruijn graph assemblers.

The simulated experiments suggested that genome length and read length influence VI; A longer genome length will produce proportionally more contigs during VI, whereas a longer read length decreases the PID range where VI occurs [Figure 4]. While longer read length improves assembly, unfortunately, platforms that produce long reads such as Oxford Nanopore and PacBio have higher error rates.^23^ Until long reads can be produced at high fidelity, researchers must continue to rely on combining long- and short-read NGS datasets, and genome polishing techniques.^23^

The large number of contigs generated due to VI may be overwhelming for most researchers, and for viral ecology studies, could lead to over-estimation of species richness for methods that use contig spectra to infer richness, such as PHACCS or CatchAll.^24, 25, 26^ This phenomenon may also impact studies differently depending on the overall goal for generating viral sequence data. For example, some researchers may only be concerned with generating a single major consensus genome, even when variants are detected in the data. This is common during outbreak responses for pathogens such as Ebola virus or Middle East respiratory syndrome coronavirus, where detection of SNPs (indicative of minor variants) is not immediately important. On the other hand, some investigations could favor distinguishing variants, such as for investigating the presence of vaccine-derived poliovirus, where a small number of SNPs may distinguish a vaccine-derived strain from a normal vaccine strain genome.^21^

The effects of VI could potentially be mitigated by running multiple assembly programs. A previous study testing bioinformatics strategies for assembling viral NGS data found that employing sequential use of de Bruijn graph and overlap-layout-consensus assemblers produced better assemblies.^15^ We speculate that this “ensemble strategy”^15^ may perform better because the multiple assemblers complement one another by having different VI PID thresholds. Future assembly approaches could also consider resolving the VI problem by possibly discriminating the major and minor variant reads first (perhaps by coverage or SNP analysis), and then assembling major and minor variant reads separately.

Since we observed VI occurring in simulated data from 2 Kb to 1 Mb genome lengths, we speculate that it may not only affect viral data but also larger draft contigs of bacteria and other microorganisms. Even though bacterial mutation rates are much lower than those of most viruses, bacterial variants are common. For environmental studies, bacterial metagenomes are known to contain many related taxa and variants ^27 28, 29, 30^, and in clinical investigations, minor bacterial variants can harbor SNPs that provide resistance against antimicrobials. This warrants future investigation into how the presence of variants may impact the assembly of other microbial datasets.

This study aimed to understand how variants affect assembly. As an initial investigation, many confounding factors were simplified for experimentation. Simulated variants studied here only depicted periodic mutations, set at regular intervals. However, in real viral data, SNPs are never evenly distributed across the genome, with zones of divergence and similarity.^31, 32^ Other important factors which influence genome assembly include sequencing error rates, presence of repetitive regions, and coverage depth. We limited our experiments to keep these factors constant in order to investigate the sole effect of VI. Through this exploration, we demonstrated that reads from related genome variants adversely affect *de novo* assembly. As NGS and *de novo* assembly have become essential for generating full-length viral genomes, future studies should investigate the combined effects of the number and relative proportion of minor variants, as well as additional assembly factors (e.g., error rates) to supplement this work.

## Methods

### Analyzing NGS and assembler usage in the virus nucleotide collection in GenBank

Viral sequence entries from the GenBank non-redundant nucleotide collection were obtained by downloading all sequences under the virus taxonomy through the end of 2017. A total of 2,338,775 GenBank entries were investigated.

The total number of viral sequences submitted annually in GenBank through December 2017 was calculated by filtering GenBank submissions by “virus,” followed by application of the following additional filtering steps: “genomic DNA/RNA” was selected and a “release date: Jan 1 through Dec 31” was applied to find the total number of viruses for a given year. A custom script was used to filter and count all documented sequencing technologies and assembly methods used for each GenBank entry.

### Creation of simulated variant genomes and reads

Simulated genomes were generated using custom scripts that randomly assign each nucleotide over a designated genome length with a weighted distribution dependent on the GC content [Supplement Figure S1]. The random genomes were then screened using NCBI BLAST to insure no similarity/identity existed to any classified organism (i.e., no BLAST hits). These simulated genomes served as the initial variant genome (variant 1). To generate the mutated variant genomes (variant 2), a custom script was used to systematically introduce evenly distributed random mutations at rates from 1 mutation in every 4 nucleotides (75% PID) to 1 mutation in every 250 nucleotides (99.6% PID), incrementing by 1 nucleotide.

Following the generation of initial and mutated variant genomes, high-quality fastq reads were generated using ART,^33^ simulating Illumina MiSeq paired-end runs at 50× coverage with 250 nt reads, DNA/RNA mean fragments size of 500, and quality score of 93. Fastq reads were combined in equal numbers for the initial and mutated variants, and used as input for subsequent *de novo* assembly experiments [Supplement Figure S1]. The same process was utilized to generate the artificial genomes, initial and mutated variant genomes, and reads for each of the experiments.

### Experiment 1: Analyzing simulated reads from variants using different *de novo* assembly programs

The simulated datasets containing reads from two variant genomes with nucleotide pairwise identity ranging from 75%–99.6% were analyzed using 10 different genome assembly programs. The *de novo* assembly algorithms used were either overlap-layout-consensus (OLC) [Cap^34^ and Mira^35, 36^], de Bruijn graph (DBG) [ABySS^37^, IDBA^38^, MetaSPAdes^39^, SOAPdenovo2^40^, SPAdes^41^, and Trinity^42^], or commercial software packages [CLC (https://www.qiagenbioinformatics.com/) and Geneious^43^] whose assembly algorithms are proprietary [Supplement Table S6]. The simulation settings for the reads were single-end reads, 250 nt read length, and 50× coverage. A total of 2,470 assemblies (247 datasets per genome × 10 assemblers) were analyzed [Supplement Figure S1a].

### Experiment 2: Simulated data by varying genome length and GC content

Artificial genomes were constructed for four genome lengths: 2 Kb, 10 Kb, 100 Kb, and 1 Mb, with varying GC content from 20%–80%, in 5% increments [Supplement Figure S1b]. Datasets derived using one poliovirus genome (NC_002058) and one coronavirus genome (NC_002645) were also included in this analysis, representing the lower and upper genome length range typical of RNA viruses. The original GC content was kept constant for the poliovirus and coronavirus genomes. For all of these genomes, simulated reads for initial and mutated variants were generated as above.

A total of 13,338 SPAdes assemblies were generated, which included 12,844 assemblies for the four artificial genomes (247 datasets per genome × 4 artificial genome lengths × 13 GC content proportions × 1 assembler) and 494 assemblies for the poliovirus and coronavirus datasets (247 datasets per genome × 2 genomes × 1 assembler) [Supplement Figure S1b]. JMP v13.0.0 (www.sas.com) was used to calculate Pearson’s correlation and Spearman’s ρ values to compare the association between percent GC levels and the number of contigs produced at each PID level. Since there was little statistical difference when comparing the contig numbers generated at varying percent GC for each of the four genome length datasets (Spearman’s ρ = 0.8299 to 0.9801, p<0.001) [Supplement Excel file], the final contig number was averaged across the 13 GC percentages at a given PID. The average contig number was used for plotting the contig assembly results vs percent PID for each simulated genome length [Figures 4a-b].

### Experiment 3: Simulated data by varying read length

Genome variants were generated as described above (“Creation of simulated variant genomes and reads”) for a genome of size 100 Kb with 50% GC; this was the starting initial variant genome. In this simulation, initial and mutation variant reads at four sequencing read lengths (50, 100, 150, and 250 nt) were created using ART. A total of 538 SPAdes assemblies were generated (47, 97, 147, and 247 datasets for the 50, 100, 150 and 250 nt read lengths, respectively) [Supplement Figure S1c].

### Evaluation of NGS datasets from clinical samples

Four datasets derived from clinical samples containing picornaviruses (one enterovirus A71 [EV-A71], one coxsackievirus B5 [CV-B5] and two parechovirus A3 [PeV-A3]) were analyzed for this experiment, as previous sequencing analysis using Geneious indicated the presence of genome variants. The datasets were analyzed using an in-house pipeline (VPipe),^18^ which performs various quality control (QC) steps and *de novo* assembly using SPAdes. The post-QC reads were considered total reads (**T**)and mapped to their respective reference genome in order to determine the major and minor variants present in each sample. Total reads which mapped with high similarity (≥99%) to the major variant were categorized as reads representing the major variant (**M**). Unbinned reads from the major variant reference recruitment were used to construct the minor variant consensus using a second round of reference recruitment, and these reads were categorized as the minor variant (**m**). Remaining reads from the previous two steps were considered background (**B**) reads.

*De novo* assembly for each of the four clinical samples was performed for the following binned NGS datasets: (1) total reads only (**T**); (2) major variants only (**M**); (3) major and minor variants only (**Mm**); and (4) major variants and background reads only (**MB**). This was repeated with three assembly programs: SPAdes, Cap3, and Geneious. The length of the longest contig produced from each assembly and the performance metric UG_50_%.^17^ were calculated to compare the results for these 48 assemblies (4 experiments × 4 viruses × 3 assemblers).

## Data Availability

Sequencing reads for the experiments conducted using clinical specimens are available through the NCBI Sequence Read Archive (SRA) accession PRJNA577924. Reads from simulated datasets (Experiments 1-3) are available upon request.

## Acknowledgements

We thank Dr. Steve Oberste for thoughtful suggestions on this work.

## Author contributions

All authors contributed to the conceptualization, data analysis, preparation, and review of this manuscript. C.J.C, R.L.M., and T.F.F.N. wrote this manuscript.

## Supplemental Information

### Analysis of viral GenBank records

#### The advent of NGS fuels viral sequencing

As of December 2017, GenBank’s non-redundant nucleotide database had grown to more than 2.3 million virus sequences, with the annual number of new sequences deposited increasing by 270% between 2007 and 2017 [Figure 1a and Supplement Table S1]. GenBank entries started incorporating information on the sequencing technology platform used in 2011. Through 2018, 144,712 viral entries (22%) had documented utilization of NGS sequencing technology, compared to 500,027 entries (77%) utilizing Sanger methods [Figure 1b and Supplement Table S1]. Illumina was the most common NGS platform used for viral sequencing, with >2-fold the number of entries compared to the next most popular NGS platform (31,000 viral entries in 2017 [Figure 1d & e]). Although NGS usage has risen tremendously, Sanger sequencing still contributed the majority of all viral sequences. This is likely because Sanger is still attractive for generating short viral sequences over genotyping windows or other informative regions. If only long sequences (≥2000 nt) are considered, NGS technologies surpassed Sanger as the dominant strategy for sequencing in 2017 [Figure 1f and Supplement Table S2].

A total of 27,217 counts of sequencing technologies were listed for the 25,344 long viral GenBank entries in 2017, as some sequences were generated using two or more sequencing technologies. NGS technologies were listed in 53.8% (14,653/27,217) of entries, versus 46.2% of entries (12,564/27,217) for Sanger. Illumina was identified as the most dominant NGS technology, accounting for 12,615/14,653 entries (86.1%) [Figure 1g and Supplement Table S2].

Multiple sequencing technologies may be used to generate viral sequence for one entry. The most common combination observed was 454 and Sanger (18,002 entries), likely due to the early emergence of the 454 technology compared to other NGS platforms [Figure 1c and Supplement Table S3]. This is followed by Illumina and Sanger (3,475), Illumina and 454 (3,299), Illumina and Ion Torrent (2,600), and Illumina and PacBio (997). Interestingly, more recently released longer-read platforms like PacBio and Oxford Nanopore tended to be paired with Illumina more frequently compared to traditional Sanger sequencing. A small number of studies even combined three or four different sequencing technologies (530 and 6 entries, respectively) [Supplement Table S4]. Some users employed a combined approach to circumvent the inherent flaws of one sequencing platform, particularly for genome finishing.^44^ For example, after NGS has been used to generate the majority of a RNA virus genome, RACE (Rapid amplification of cDNA ends) is typically performed with Sanger to obtain the 5’ or 3’ termini.^45, 46^

#### *De novo* assembly plays a major role in analyzing long viral sequences

We analyzed the assembly methods used for GenBank entries of long sequences (≥2000 nt) from 2012 to 2017 when NGS usage become relevant [Figure 1h & i and Supplement Table S5]. The number of programs used to assemble viral sequences has steadily increased over time (a >2-fold increase from 2012-2017). With new sequencing technologies emerging and computational power continually improving, the development of new and better assembly programs always follows suite. The use of specifically-designed *de novo* assembly programs (ABySS, BWA, Canu, Cap3, IDBA, MIRA, Newbler, SOAPdenovo, SPAdes, Trinity, and Velvet) has increased from less than 1% of viral sequence entries in 2012, to 20% of all viral sequence entries in 2017. A similar increase was observed for reference-mapping software (i.e., Bowtie and Bowtie2), from 0.03% in 2012 to 6.5% in 2017. Multifunctional programs that offer both assembly options, including CLC Genomics Workbench (CLC), DNA Baser, DNASTAR, Geneious, and Sequencher, were by far the most popular option for the years 2013-2017. However, since these commercial software packages can perform both *de novo* and reference-mapping assembly, the exact sequence assembly strategy used for these records is unknown, and thus the contributions of both *de novo* assembly and reference recruitment are likely underestimated.

---

**Supplement Figure S1.**
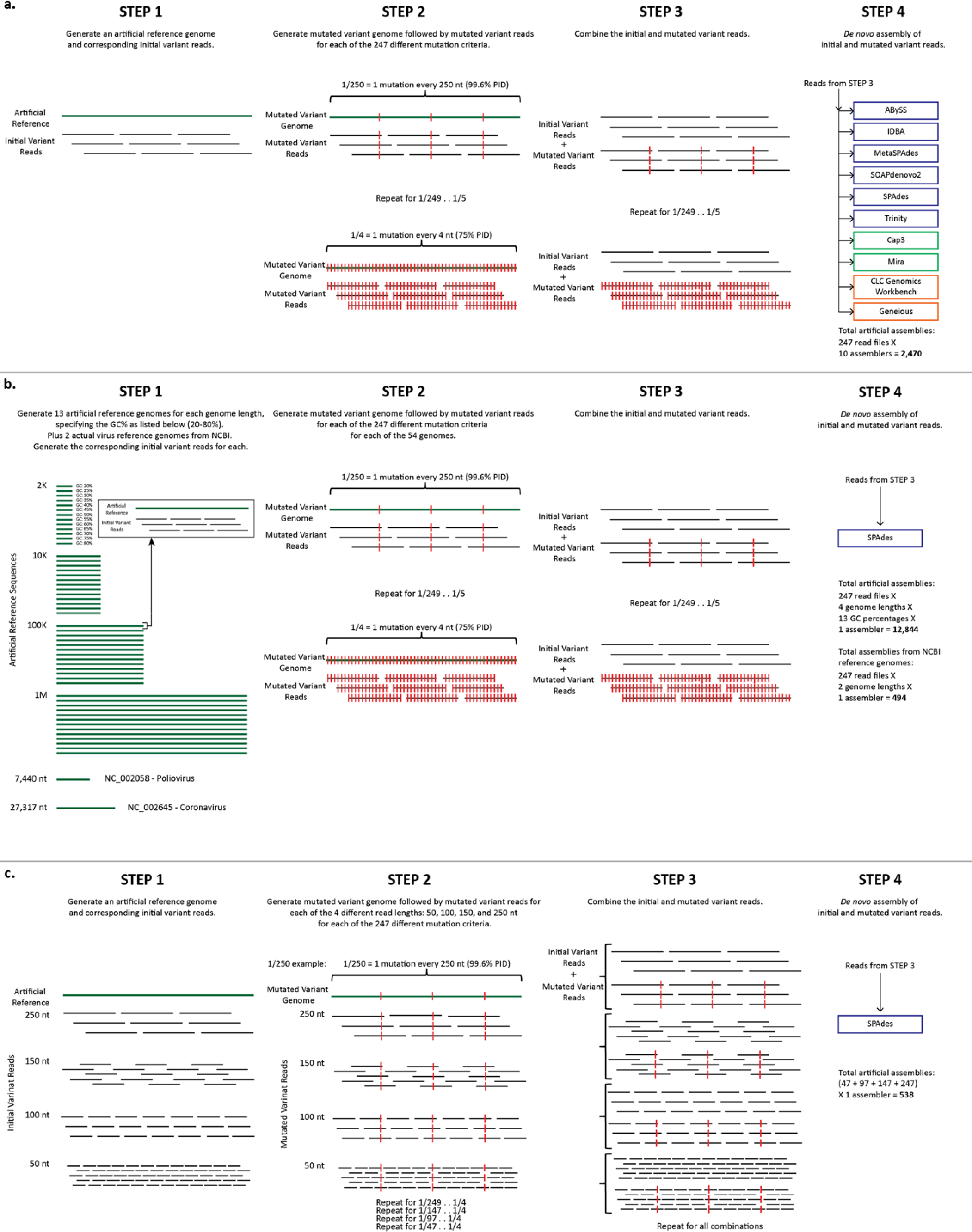
Workflow diagrams of simulated data from data creation through *de novo* assembly. **(a) Comparison of assemblers.** First, an artificial reference genome and corresponding initial variant reads were created with the following constraints: (1) reference genome length: 100K; (2) GC% of reference genome: 50%; (3) read length: 250 nt; and (4) coverage: 50×. Second, an artificial mutated variant genome and corresponding mutated variant reads were created 247 times, each with a differing pairwise percent identity ranging from 1 mutation every 4 nucleotides (75% PID) to 1 mutation in every 250 nucleotides (99.6% PID). The initial and mutated variants were then combined and used as input for 10 different *de novo* assemblers with varying underlying algorithms. A total of 2,470 assemblies were performed. **(b) Comparison of genome length and GC%.** First, 13 artificial reference genomes and corresponding initial variant reads were created for four different genome lengths (2Kb, 10Kb, 100Kb, and 1Mb), each specifying a different GC% ranging from 20%–80%. In addition, two actual virus reference genomes from NCBI were included, NC_002058 and NC_002645, with genome lengths of 7,440 nt and 27,317 nt, respectively. Read lengths of 250 nt with a coverage of 50X were used for all genomes. Second, an artificial mutated variant genome and corresponding mutated variant reads were created 247 time, each with a differing pairwise percent identity ranging from 1 mutation every 4 nucleotides (75% PID) to 1 mutation in every 250 nucleotides (99.6% PID). The initial and mutated variants were then combined for each and used as input for the SPAdes *de novo* assembler. A total of 13,338 assemblies were performed. **(c) Comparison of read length.** First, an artificial reference genome and corresponding initial variant reads were created with the following constraints: (1) reference genome length: 100K; (2) GC% of reference genome: 50%; (3) read lengths: 50 nt, 100 nt, 150 nt, or 250 nt; and (4) coverage: 50×. Second, an artificial mutated variant genome and corresponding mutated variant reads were created, each with a differing pairwise percent identity ranging from 1 mutation every 4 nucleotides (75% PID) up to 1 mutation in every 250 nucleotides (99.6% PID). The initial and mutated variants created for each of the four read lengths were then grouped by read length size and used as input for SPAdes *de novo* assembler. A total of 538 assemblies were performed.

**Supplement Figure S2.**
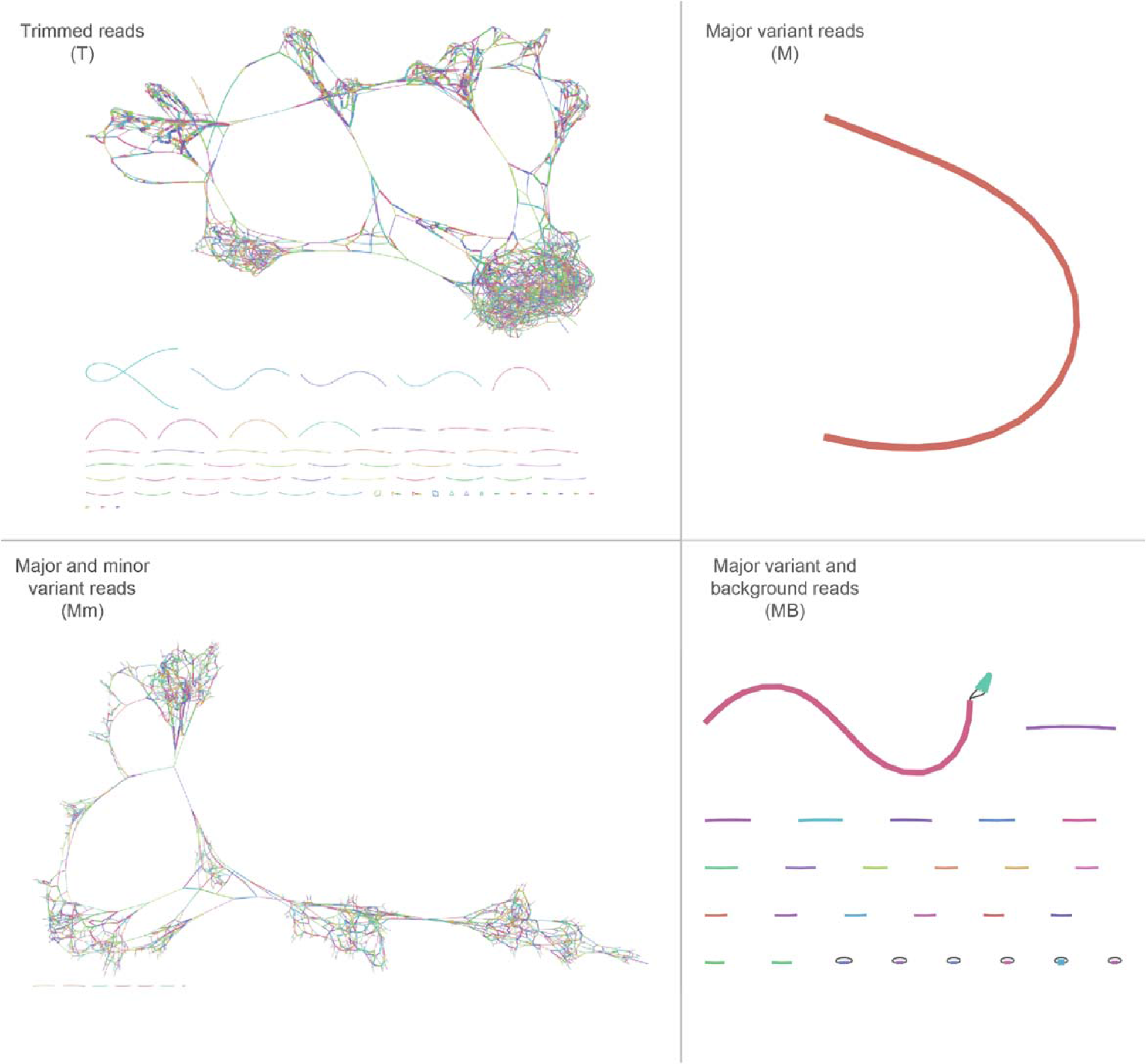
Analysis of the final contig assembly graphs for a clinical sample containing enterovirus A71 (EV-A71) variants using Bandage. Based on the four assemblies in Figure 5, Bandage was used to display the contig graphs from each SPAdes output. The visualizations for T, Mm, and MB show the effects of variant interference, while M shows the ideal assembly.

**Supplement Table S1.** Total counts from NCBI’s GenBank non-redundant nucleotide database. **†** *Total count* is the combination of all sequencing technologies listed for each entry plus the total number of entries with sequencing technology omitted. This number is higher than the *Total # of viral entries in GenBank* because it accounts for all entries with multiple sequencing technologies listed. Sequencing Technology, Seq. Tech.; Oxford Nanopore, Oxford NP; Pacific Biosciences, PacBio

| Year | Total # of viral<br>entries in GenBank | Total<br>count† | Total<br>omitted | Total # of entries with<br>at least one Seq. Tech. | Sequencing Technology Break |  |  |  |  |
| --- | --- | --- | --- | --- | --- | --- | --- | --- | --- |
|  |  |  |  |  | Sanger | 454 | Illumina | IonTorrent | Oxford N |
| 2017 | 238367 | 243849 | 108021 | 135828 | 85194 | 15999 | 31279 | 2130 | 46 |
| 2016 | 235477 | 237569 | 107090 | 130479 | 102837 | 2971 | 22185 | 2111 | 119 |
| 2015 | 197440 | 211177 | 71148 | 140029 | 102440 | 15517 | 17625 | 3048 |  |
| 2014 | 158579 | 163092 | 66217 | 96875 | 81515 | 5452 | 7399 | 2345 |  |
| 2013 | 198540 | 202232 | 108365 | 93867 | 84527 | 5243 | 2474 | 758 |  |
| 2012 | 172850 | 173324 | 126821 | 46503 | 43509 | 1194 | 277 | 403 |  |
| 2011 | 181315 | 181319 | 176355 | 4964 | 5 | 4811 | 147 |  |  |
| 2010 | 131962 | 131962 | 131960 | 2 | 2 |  |  |  |  |
| 2009 | 213549 | 213549 | 213549 |  |  |  |  |  |  |
| 2008 | 109265 | 109265 | 109265 |  |  |  |  |  |  |
| 2007 | 88996 | 88996 | 88996 |  |  |  |  |  |  |
| 2006 | 94444 | 94444 | 94444 |  |  |  |  |  |  |
| 2005 | 58245 | 58245 | 58245 |  |  |  |  |  |  |
| 2004 | 53841 | 53842 | 53834 | 8 | 1 | 4 | 2 |  |  |
| 2003 | 38578 | 38578 | 38576 | 2 | 2 |  |  |  |  |
| 2002 | 33412 | 33412 | 33412 |  |  |  |  |  |  |
| 2001 | 28305 | 28305 | 28304 | 1 |  | 1 |  |  |  |
| 2000 | 26871 | 26871 | 26871 |  |  |  |  |  |  |
| 1999 | 17266 | 17266 | 17266 |  |  |  |  |  |  |
| 1998 | 13840 | 13840 | 13840 |  |  |  |  |  |  |
| 1997 | 12378 | 12378 | 12378 |  |  |  |  |  |  |
| 1996 | 8988 | 8988 | 8987 | 1 |  |  | 1 |  |  |
| 1995 | 7475 | 7475 | 7475 |  |  |  |  |  |  |
| 1994 | 5449 | 5449 | 5449 |  |  |  |  |  |  |
| 1993 | 9185 | 9185 | 9184 | 1 |  |  | 1 |  |  |
| 1992 | 1754 | 1754 | 1754 |  |  |  |  |  |  |
| 1991 | 725 | 725 | 725 |  |  |  |  |  |  |
| 1990 | 364 | 364 | 363 | 1 |  |  | 1 |  |  |
| 1989 | 424 | 424 | 424 |  |  |  |  |  |  |
| 1988 | 269 | 269 | 269 |  |  |  |  |  |  |
| 1987 | 159 | 159 | 159 |  |  |  |  |  |  |
| 1986 | 114 | 114 | 114 |  |  |  |  |  |  |
| 1985 | 130 | 130 | 130 |  |  |  |  |  |  |
| 1984 | 19 | 19 | 19 |  |  |  |  |  |  |
| 1983 | 92 | 92 | 92 |  |  |  |  |  |  |
| 1982 | 108 | 108 | 108 |  |  |  |  |  |  |
| TOTALS | 2338775 | 2368770 | 1720209 | 648561 | 500032 | 51192 | 81391 | 10795 | 165 |

**Supplement Table S2.** Total count of sequencing technologies for sequences >2000 nt in the NCBI GenBank non-redundant nucleotide database for years 2012–2017. These numbers were found with the following search criteria: “viruses,” “genomic RNA/DNA,” “GenBank (No RefSeq),” length: 2000 to 2000000, release date: 1/1/201X to 12/31/201X, and “sequencing technology” in any field. Oxford Nanopore, Oxford NP; Pacific Biosciences, PacBio

| NGS<br>Platforms | Year |  |  |  |  |  |
| --- | --- | --- | --- | --- | --- | --- |
|  | 2017 | 2016 | 2015 | 2014 | 2013 | 2012 |
| **454** | 1029 | 634 | 4987 | 1531 | 1642 | 376 |
| **Sanger** | 12564 | 13571 | 20216 | 14294 | 13646 | 10847 |
| **Illumina** | 12615 | 12629 | 4121 | 4414 | 1266 | 230 |
| **PacBio** | 17 | 67 | 1 | 12 | 1 | 0 |
| **IonTorrent** | 923 | 1342 | 1217 | 1131 | 408 | 171 |
| **Oxford NP** | 46 | 119 | 0 | 0 | 0 | 0 |
| **SOLiD** | 8 | 0 | 0 | 13 | 29 | 1 |
| **Other** | 15 | 4 | 1 | 5 | 10 | 41 |
| **TOTALS** | **27217** | **28366** | **30543** | **21400** | **17002** | **11666** |

**Supplement Table S3.** Total counts from NCBI’s GenBank non-redundant nucleotide database with multiple sequencing technologies listed per entry. Blank fields indicate absence of entries for the corresponding category. Sequencing Technologies, Seq. Techs.

| Year | Total # of entries with<br>two Seq. Techs. | Total # of entries with<br>three Seq. Techs. | Total # of entries with<br>four Seq. Techs. |
| --- | --- | --- | --- |
| 2017 | 5468 | 7 |  |
| 2016 | 2008 | 42 |  |
| 2015 | 13156 | 283 | 5 |
| 2014 | 4457 | 28 |  |
| 2013 | 3409 | 140 | 1 |
| 2012 | 414 | 30 |  |
| 2011 | 4 |  |  |
| 2010 |  |  |  |
| 2009 |  |  |  |
| 2008 |  |  |  |
| 2007 |  |  |  |
| 2006 |  |  |  |
| 2005 |  |  |  |
| 2004 | 1 |  |  |
| 2003 |  |  |  |
| 2002 |  |  |  |
| 2001 |  |  |  |
| 2000 |  |  |  |
| 1999 |  |  |  |
| 1998 |  |  |  |
| 1997 |  |  |  |
| 1996 |  |  |  |
| 1995 |  |  |  |
| 1994 |  |  |  |
| 1993 |  |  |  |
| 1992 |  |  |  |
| 1991 |  |  |  |
| 1990 |  |  |  |
| 1989 |  |  |  |
| 1988 |  |  |  |
| 1987 |  |  |  |
| 1986 |  |  |  |
| 1985 |  |  |  |
| 1984 |  |  |  |
| 1983 |  |  |  |
| 1982 |  |  |  |
| **TOTALS** | **28917** | **530** | **6** |

**Supplement Table S4.**
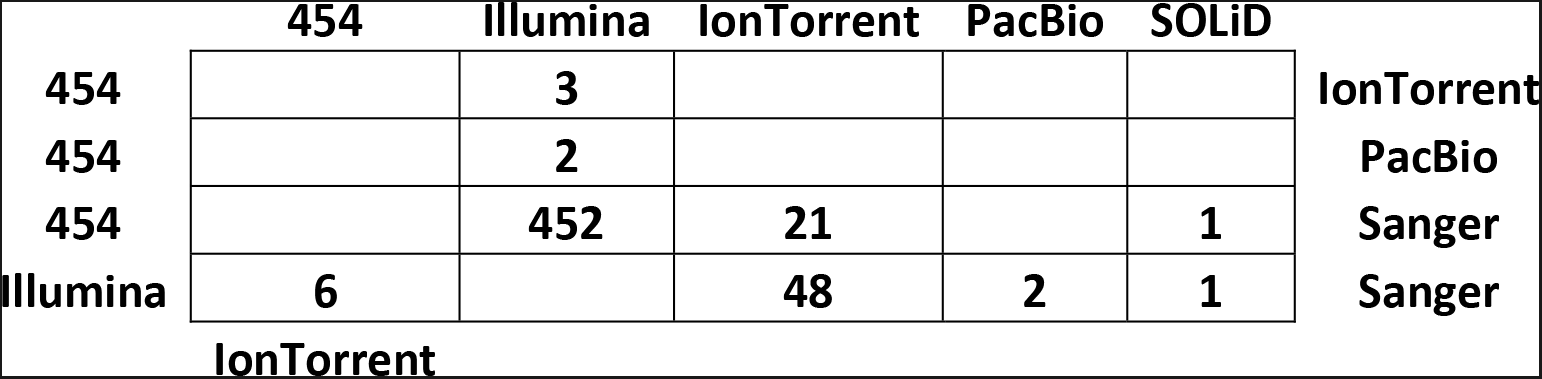
Total counts from NCBI’s GenBank non-redundant nucleotide database of all entries with three and four sequencing technologies listed. For example, there are a total of 6 entries in GenBank that have the following sequencing technologies listed: 454, Illumina, Ion Torrent, and Sanger for one sequence entry. Pacific Biosciences, PacBio

**Supplement Table S5.** Total count of assembly programs used to generate sequences >2000 nt in the NCBI GenBank non-redundant nucleotide database. These numbers were found with the following search criteria: “viruses,” “genomic RNA/DNA,” “GenBank (No RefSeq),” length: 2000 to 2000000, release date: 1/1/201X to 12/31/201X, and ‘”sequencing technology” in any field; the assembly method was then parsed out.

| Assembly<br>Methods | Year |  |  |  |  |  |
| --- | --- | --- | --- | --- | --- | --- |
|  | 2017 | 2016 | 2015 | 2014 | 2013 | 2012 |
| **ABYSS** | 522 | 155 | 100 | 66 | 56 | 0 |
| **Bowtie** | 40 | 868 | 33 | 527 | 5 | 4 |
| **Bowtie2** | 1682 | 128 | 787 | 9 | 51 | 0 |
| **BWA** | 671 | 294 | 281 | 440 | 148 | 1 |
| **Canu** | 3 | 0 | 0 | 0 | 0 | 0 |
| **Cap3** | 59 | 34 | 55 | 288 | 10 | 0 |
| **CLC** | 3404 | 5139 | 1948 | 2186 | 1172 | 381 |
| **DNA Baser** | 838 | 326 | 247 | 261 | 27 | 9 |
| **DNASTAR** | 4030 | 3191 | 6897 | 3175 | 3101 | 530 |
| **Geneious** | 3636 | 2633 | 4767 | 588 | 504 | 79 |
| **IDBA** | 28 | 11 | 729 | 22 | 2 | 0 |
| **MIRA** | 446 | 406 | 70 | 140 | 24 | 14 |
| **Newbler** | 176 | 183 | 703 | 336 | 435 | 60 |
| **Sequencher** | 3243 | 2154 | 2572 | 5727 | 7927 | 3462 |
| **SOAPdenovo** | 258 | 67 | 105 | 24 | 9 | 1 |
| **SPAdes** | 792 | 1632 | 89 | 266 | 0 | 0 |
| **Trinity** | 2162 | 4576 | 301 | 509 | 4 | 0 |
| **Velvet** | 161 | 107 | 338 | 341 | 144 | 32 |
| **Other** | 4190 | 6220 | 5810 | 5437 | 3179 | 6950 |
| **TOTALS** | **26341** | **28124** | **25832** | **20342** | **16798** | **11523** |

**Supplement Table S6.** Supplement Table S6. The 10 *de novo* assemblers used for analysis of the simulated data, as categorized by their underlying assembly algorithms. de Bruijn graph, DBG; overlap-layout-739 consensus, OLC.

| DBG |  | OLC |  | Proprietary Algorithm |  |
| --- | --- | --- | --- | --- | --- |
| Program | Version | Program | Version | Program | Version |
| ABYSS | 2.0.2 | Cap | 3 | CLC Genomic Workbench | 11 |
| IDBA | 1.1.3 | Mira | 4.0.2 | Geneious | 10.2.3 |
| MetaSPAdes | 3.9.0 |  |  |  |  |
| SOAPdenovo2 | r240 |  |  |  |  |
| SPAdes | 3.9.0 |  |  |  |  |
| Trinity | 2.1.1 |  |  |  |  |

